# Investigations into genetic control of six spike traits with a focus on breeding for terminal heat stress tolerance in common wheat

**DOI:** 10.1101/2025.06.30.662399

**Authors:** Sourabh Kumar, Sachin Kumar, Vivudh Pratap Singh, Hemant Sharma, Kanwardeep Singh Rawale, Sunil Kumar Bhatt, Ramanathan Vairamani, Kulvinder Singh Gill, Harindra Singh Balyan

**Affiliations:** Department of Genetics and Plant Breeding, Chaudhary Charan Singh University, Meerut, Uttar Pradesh, India; Department of Botany, Chaudhary Charan Singh University, Meerut, Uttar Pradesh, India; Geneshifters, LLC, Pullman, Washington, USA; Research and Development Division, JK Agri-Genetics Limited, Hyderabad, Telangana, India; Rallis India Limited, Mumbai, Maharashtra, India; Department of Crop and Soil Sciences, Washington State University, Pullman, Washington, USA

**Keywords:** Wheat, terminal heat stress, SNPs, quantitative trait loci, candidate genes, KASP marker, gene-based functional markers

## Abstract

To unravel the genetic architecture of six spike traits under heat stress, we used a doubled-haploid (DH) population (177 lines), developed from a cross between a heat-sensitive cultivar (PBW343) and a heat-tolerant genotype (KSG1203). This DH population and the two parents were phenotyped under timely, late, and very late sown conditions across 15 environments over three years and two locations. Best linear unbiased estimates and a high-density genetic map (5,710 SNP markers) were used for QTL mapping. A total of 51 QTL were detected in timely (17), late (10), and very late (18) sown conditions, with six common across environments with phenotypic variation explained ranging from 7.1% to 23.6%. A set of 14 stable, major QTL were validated in high-yielding DH lines and are recommended for marker-assisted recurrent selection. Several QTL co-localized with known genes responsible for important traits including grain yield (*TaGW2-B1*, *PI1-1B*). Seventy heat-responsive candidate genes associated with QTL were identified, encoding 33 proteins. A KASP marker for floret fertility QTL (*QFf.ccsu-3A*), and gene-based SSR markers for five key genes were developed and validated alongside Indels and SNPs. The generated genomic resources could be used in future studies and to breed heat-tolerant, high-yielding wheat varieties and germplasm.

**Highlight:** Identified and validated stable QTL for spike traits across 15 environments, promising candidate genes for thermotolerance and spike traits, novel KASP and gene-based markers, providing genomic resources for breeding high-yielding, heat-tolerant wheat.

**Graphical Abstract:** 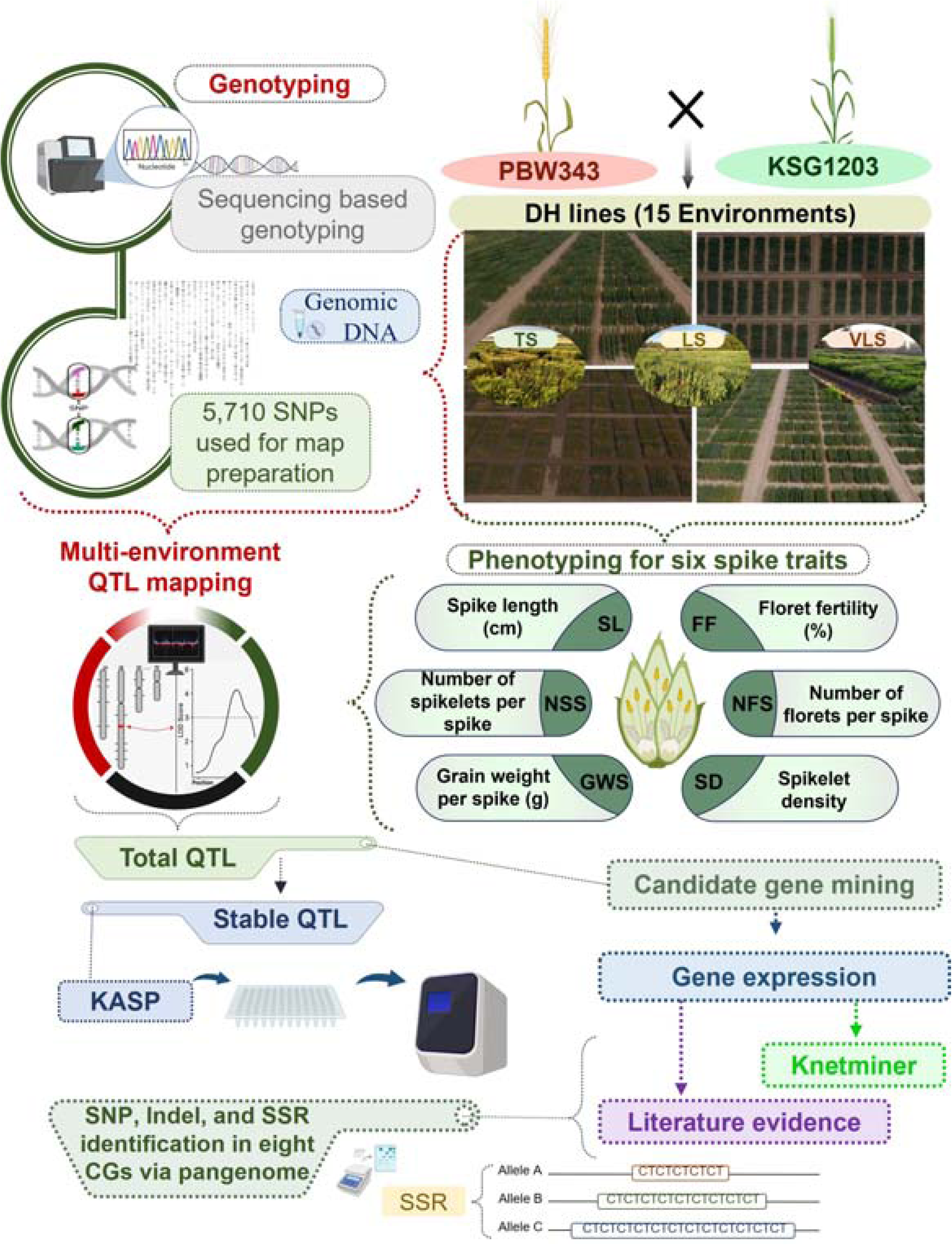

## Introduction

Wheat (*Triticum aestivum* L.) is a globally cultivated staple crop, with production estimated at 784.91 million metric tons (Mt) in 2023–24, down by 4.26 Mt from the previous year (https://tinyurl.com/statisticsreportwheat). About 25% of global production is traded internationally, and nearly 55% of the global population relies on wheat, which supplies around 20% of daily carbohydrate and protein intake, underscoring its importance in nutrition and food security (Langridge *et al.,* 2022; Bhatt *et al.,* 2024). To meet the needs of a projected global population of 9.7 billion by 2050, particularly in Asia, wheat production must increase to ∼858 Mt, a 60% rise from 2010 levels (Hickey *et al.,* 2019; Rosegrant and Agcaoili, 2010). This requires at least 0.7% annual genetic yield improvement (Ray *et al.,* 2013; https://www.cimmyt.org/work/wheat-research/).

Climate change poses major threats to this goal, introducing new abiotic and biotic stresses. Globally, ∼50% of wheat areas are affected by heat stress, while 11% suffer from consistent water deficits. Combined, these stresses contribute to 40% of annual yield variability (Zampieri *et al.,* 2017). In the Indo-Gangetic Plains (IGP) of India, a critical wheat-producing region, rising minimum temperatures have been linked to declining potential yields (Pathak *et al.,* 2003). Projections suggest that heat stress could reduce Indian wheat production by 10–40% by 2080–2100 (Boraiah *et al.,* 2023). Since 1980, global temperatures have risen by 0.8 °C, with two-thirds of the increase occurring after 1975 at a rate of 0.15–0.20 °C per decade (Lorenz *et al.,* 2019). The IPCC projects a 0.7–2.0 °C increase in India by the 2030s and 3.3–4.8 °C by the late 21^st^ century (IPCC, 2014). During the 20^th^ century, wheat-season temperatures in India rose by 0.04 °C per decade (Chauhan *et al.,* 2014). In South Asia’s rice-wheat systems, including India, heat stress is already a concern, with daily averages exceeding 17.5 °C in the coolest month (Fischer and Byerlee, 1991). It is feared that IGP may become unsuitable for wheat cultivation by 2050, with each 1 °C increase in temperature potentially reducing yields by 4–5 Mt (Boraiah *et al.,* 2023). Even a 1–2 °C rise could cause a 7–21% drop in production (Kristensen *et al.,* 2011; Liu *et al.,* 2016; Zaveri and Lobell, 2019).

All growth stages of wheat are vulnerable to heat, but terminal heat stress, occurring during reproduction and maturity, is especially damaging, leading to substantial yield losses (Parent *et al.,* 2017; Jacott and Boden, 2020; Farhad *et al.,* 2023; Mehmood *et al.,* 2025). The PEGASUS crop model estimates that heat stress during anthesis may halve projected yield gains (Derying *et al.,* 2014). In Punjab, India, recent heatwaves at grain filling stage triggered premature ripening, shrivelling, and yellowing of grains, resulting in a 25% yield loss (Bal *et al.,* 2022). Terminal heat stress causes numerous physiological disruptions such as pollen sterility, dehydration, reduced CO assimilation, increased photorespiration, and lower pollen viability (Farooq *et al.,* 2011; Jagadish, 2020). Heat stress during reproduction hampers gametophyte development, reduces pollen germination by 39.9%, seed number by 23%, and seed weight by 34.6% on main/primary spikes (Bheemanahalli *et al.,* 2019; Qian *et al.,* 2025; Mehmood *et al.,* 2025). It impairs inflorescence and spikelet development, reduces grain size and weight, delays spikelet initiation, and often leads to infertility (Jacott and Boden, 2020; Qian *et al.,* 2025). Just one hot hour (>28 °C) before flowering can reduce grain count by 0.25 grains per spike and 281 grains/m². Post-flowering exposure to >32 °C can lower grain weight by 0.26 mg and total yield by 244 g/m² (Ullah *et al.,* 2024). Grain-filling duration may be shortened by 45–60%, affecting grain size, weight, and quality through altered enzymatic function and source-sink relationships (Sehgal *et al.,* 2018; Abdelrahman *et al.,* 2020a; Khanzada *et al.,* 2025). High night temperatures also reduce agronomic performance, with trait reductions up to 35% in tolerant genotypes and up to 75% in susceptible ones (Kahlon *et al.,* 2024). Overall, heat stress could reduce field-level grain production by as much as 60% (Zampieri *et al.,* 2017).

The rice-wheat and sugarcane-wheat cropping systems in IGP often delay wheat sowing, subjecting crops to 32–38 °C during grain development—well above the optimal 22–25 °C. Even timely sowing cannot always prevent heat exposure due to shifting climate patterns. In fact, a recent study found only one-third of genotypes tested showed good adaptation, with stable phenotypes observed in only 26% of environments under a 0.26 °C warming trend (Xiong *et al.,* 2024). Thus, breeding heat-resilient, high-yielding wheat is an urgent priority. This requires deciphering the genetic basis of heat-responsive traits such as those linked to spike development and fertility, enabling breeders to use marker-assisted selection (MAS). Though many QTL related to heat tolerance have been reported, their application in breeding remains limited (Singh et al. 2022; http://www.wheatqtldb.net/).

Several QTL studies in IGP and beyond have used population-specific genetic maps to dissect heat-responsive traits (Kumar *et al.,* 2010; Paliwal *et al.,* 2012; Tiwari *et al.,* 2013; Sharma *et al.,* 2016; Bhusal *et al.,* 2017, 2018; Raveendran *et al.,* 2020; Pankaj *et al.,* 2022; Manjunath *et al.,* 2024; Singh *et al.,* 2024). Genetic maps provide chromosomal locations and effect sizes of QTL, aiding MAS. However, physical maps offer finer resolution, enabling comparative analysis and functional validation (Semagn *et al.,* 2021; Kumar *et al.,* 2024). Advances like the IWGSC RefSeq v2.0 (http://wheat-urgi.versailles.inra.fr/) have improved marker density and accuracy across chromosomes, enhancing QTL alignment across studies and facilitating candidate gene discovery.

The present study focuses on identifying QTL for terminal heat stress tolerance in wheat by analyzing six critical spike traits. A doubled haploid (DH) population was developed from a cross between a widely grown mega-variety in Southeast Asia and a heat-tolerant wheat selection (unpublished data, Kulvinder Gill et al.). The population, comprising 177 lines, was phenotyped in 15 field trials over three years across two IGP locations under timely, late, and very late sowing conditions. Sequencing-based genotyping (SBG) was used to generate SNP marker data, which, along with phenotypic data, facilitated QTL mapping. Identified QTL were localized on the IWGSC RefSeq v2.0 physical map, and candidate genes within QTL regions were explored via *in silico* expression analysis. A user-friendly KASP marker associated with a major, stable QTL for floret fertility was developed, along with SSR markers for five candidate genes developed with the help of pangenome. The study also proposes the development of other pangenome aided gene-specific markers based on Indels and SNPs discovered during the present study, offering essential genomic tools for breeding terminal heat-tolerant wheat varieties.

## Materials and Methods

### Genetic materials

Present study used a DH mapping population of common wheat, consisting of 177 lines. This population was generated from a cross between PBW343, a heat-sensitive mega variety common in Southeast Asia, and KSG1203, a selection from the Egyptian cultivar Giza168 and known for its tolerance to heat stress (Kumar *et al.,* 2024). The DH lines were created using a wheat × maize hybridization system, following the method described by Laurie and Bennett (1986).

### Field trials and temperature stress

A total of 180 wheat genotypes were assessed, consisting of 177 DH lines, two parental genotypes (PBW343 and KSG1203), and HD2967, which served as a filler. The study employed an alpha-lattice design with three replications and was conducted at two locations: Meerut (latitude 28° 58’ N, longitude 77° 44’ E) and Lucknow (latitude 26° 53’ N, longitude 81° 5’ E). The trials at Meerut spanned three planting periods: November 11-20 for timely sown (TS), December 17-23 for late sown (LS), and January 12-23 for very late sown (VLS), and at Lucknow: November 21-23 for TS, December 14-24 for LS, and January 12-19 for VLS. These two sites, located about 500 km apart, are part of the IGP region.

At Meerut, trials were conducted over three consecutive growing seasons (2017-18, 2018-19, and 2019-20) at the Agriculture Research Farm, Department of Genetics and Plant Breeding, Chaudhary Charan Singh University, Meerut. At Lucknow, trials were conducted at the following two different research farms: (i) JK Agri-Genetics Limited, Hyderabad, Telangana in 2017-18, and (ii) the Regional Station of Rallis India Limited, Wadala, Mumbai, in 2018-19. Altogether, the experimental setup spanned 15 environments (E1-E15, Table 1).

**Table 1.** Details of 15 individual environments (E1 to E15) and the nine pooled environments (TS1-TS3, LS1-LS3, and VLS1-VLS3) used for statistical analysis in the present study.

| Sowing condition | Meerut |  |  |  | Lucknow |  |  | Overall Pooled** | Total number of pooled environment |
| --- | --- | --- | --- | --- | --- | --- | --- | --- | --- |
|  | 2018 | 2019 | 2020 | Pooled* | 2018 | 2019 | Pooled* |  |  |
| TS | E1 | E7 | E13 | **TS1** | E2 | E8 | **TS2** | **TS3** | 3 (TS1, TS2, TS3) |
| LS | E3 | E9 | E14 | **LS1** | E4 | E10 | **LS2** | **LS3** | 3 (LS1, LS2, LS3) |
| VLS | E5 | E11 | E15 | **VLS1** | E6 | E12 | **VLS2** | **VLS3** | 3 (VLS1, VLS2, VLS3) |

TS: timely sown, LS: late sown, VLS: very late sown; \*Data pooled over the years separately at Meerut and Lucknow locations; \*\*Data pooled across the locations and years

The alpha-lattice experiments were conducted with three replicates. Each replicate was divided into 18 blocks, with each block containing 10 plots, each assigned to different genotypes. The plots were 2.5 meters in length and consisted of four rows spaced 20 cm apart. Twenty grams of seed per plot was used for planting, corresponding to a seeding rate of 120 kg per hectare. Irrigation was provided four to six times as needed. Fertilizers were applied at standard rates: 120 kg nitrogen (N) as urea (46% N), 60 kg phosphorus (P) as single super phosphate (16% P), and 40 kg potassium (K) as murate of potash (60% K). Full doses of phosphorus and potassium were applied at sowing, along with one-third of the nitrogen and the rest split between the first and second irrigations. To control rust, the wheat trials were treated twice with the fungicide Tilt (propiconazole), once before anthesis and once afterward. Temperature data, including daily minimum and maximum values, were collected from nearby weather stations in Meerut and Lucknow for each cropping season.

### Trait phenotyping

Phenotypic data were gathered for six spike traits across 15 field trials: (i) Spike length (SL), (ii) Number of spikelets per spike (NSS), (iii) Spikelet density (SD), (iv) Number of florets per spike (NFS), (v) Floret fertility (FF), and (vi) Grain weight per spike (GWS). Detailed methods for measuring these traits are provided in Supplementary Table S1.

### Statistical analysis

A combined analysis of variance (ANOVA) was performed using phenotypic data from 15 different environments (E1-E15), which varied in sowing conditions, locations, and years (Table 1). The analysis was conducted with R software packages like ‘metan’ (https://CRAN.R-project.org/package=metan), ‘lme4’ (https://cran.r-project.org/web/packages/lme4/index.html), and ‘agricolae’ (https://cran.r-project.org/web/packages/agricolae/index.html). Descriptive statistics and broad-sense heritability (H²) for each trait in TS, LS and VLS trials were calculated using the R packages ‘variability’ (https://CRAN.R-project.org/package=variability) and ‘lme4’ involving the following formula:

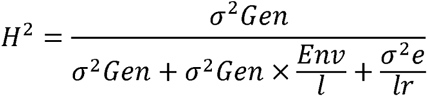

where σ^2^Gen = genotypic variance, σ^2^Gen × Env = genotype × environment interaction variance, σ^2^e = error variance, l = number of environments and r = number of replications.

The best linear unbiased estimates (BLUEs) for all six spike traits in the DH population were derived from phenotypic data collected across the TS, LS, and VLS environments. BLUEs were calculated separately for the TS, LS, and VLS environments over multiple years at the two locations, as well as combined across both locations. Table 1 provides the specifics of the 15 individual environments and nine pooled environments that resulted from this process. The pooled BLUEs used for further analysis were generated following the standard procedure in METAR software (Alvarado *et al.,* 2020), using the following model:

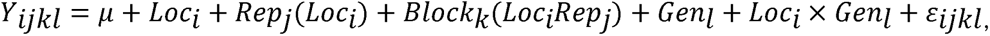

where Yijkl is the response value of observed trait, µ the overall mean, Loc_i_ is the location effect, Rep_j_ is the replicate effect (j = 1, 2, 3, …, n), Block is the block effect, Gen_l_ is the treatment fixed effect (l = 1, 2, 3, …, n), and εijkl is the error term.

Ridgeline plots displaying the BLUEs for six spike traits were generated using the R package ‘ggridges’ (https://cran.r-project.org/web/packages/ggridges/vignettes/introduction.html). Pearson’s correlation coefficient analyses were conducted among the six spike traits across three sowing environments (TS, LS, and VLS) by utilizing the pooled BLUE values over multiple years and locations, with the help of the R package ‘metan’ (https://CRAN.R-project.org/package=metan). The path coefficient analysis was conducted following the method proposed by Dewey and Lu in 1959. It utilized phenotypic correlation coefficients to determine the direct and indirect effects of various spike traits on the GWS. This analysis was performed using the following equation implemented in the R software package ‘variability’ (https://CRAN.R-project.org/package=variability).

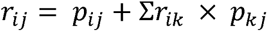

where, *r_ij_* is the mutual association between the independent trait (i) and the dependent trait (j) as the measure by the correlation coefficient, *p_ij_* is the component of direct effects of the independent trait (i) on the dependent variable (j), and *r_ik_p_kj_* is the assumption of components of the direct effect of a given independent trait via all other independent traits.

The residual effect was obtained as per the following formula:

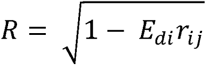

where, di is direct effect of the trait, *r_ij_* is correlation coefficient of ith trait with jth trait.

### DNA extraction, genotyping and SNP calling

The details of the procedures followed for genomic DNA extraction from seedlings of the two parents and 177 DH lines and for the determination of DNA quality are described in our recent study (Kumar *et al.,* 2024). The good quality DNA was subjected to sequencing based genotyping (SBG) by Geneshifters, Pullman, USA using their proprietary method of SNP genotyping (Kahlon *et al.,* 2024). Sequence for gene fraction were enriched to obtain 23-29% wheat genic fraction. Briefly, ∼100 ng DNA from each plant was used for restriction digestion using *PstI* and *MspI* for the preparation of library followed by barcode ligation of the fragmented DNA and subsequent pooling. These pooled samples were amplified using PCR, cleaned and fragments of ∼200 bp were selected and eluted as described in Kumar *et al.,* (2024). After size selection and quantification, the library was subjected to sequencing using Illumina Sequencer. Based on the bar codes, the raw sequence data was assigned to individual samples. Raw sequencing data was uniquely mapped to the high-confidence gene annotation downloaded from the IWGSC webserver using the bowtie2 program. Duplicate mapped reads were filtered out, and SNP calling was done using samtools and Vcftools programs (Danecek *et al.,* 2011) with a minimum sequencing depth of 10 reads per SNP. SNP call data were sorted based on the wheat physical reference genome and were converted to ‘A’ or ‘B’ format using the standalone tassel package (Bradbury *et al.,* 2007).

### Construction of genetic map

Details of the construction of the genetic map are presented in our recent study (Kumar *et al.,* 2024). In summary, 26,213 SNP markers showing polymorphism between the parental genotypes of the mapping population were identified; out of these, 20,503 markers showing either segregation distortion or >20% missing data was excluded from further analysis leaving 5,710 high-quality SNP markers. These markers were used in linkage analysis and genetic map construction using MSTMap software (Wu *et al.,* 2008). The obtained genetic map was refined using JoinMap v4.0 (Ooijen, 2006), with a logarithm of odds (LOD) threshold of 5 and a recombination frequency limit of 0.30. The recombination fractions were converted to centimorgan (cM) distances using the Kosambi mapping function (Kosambi, 1944). Linkage groups, excluding markers with unknown chromosomes (ChrUn), were assigned to specific wheat chromosomes based on the Chinese Spring reference genome (IWGSC RefSeq v2.0). The final maps for each linkage group were generated using MapChart v2.2 (Voorrips, 2002).

### Preparation of physical map of QTL

The physical map of QTL was prepared essentially in the same manner as described in our recent study by Kumar *et al.,* (2024). Briefly, the physical positions of the sequences of the SBG tags for the 5,710 SNP markers were determined following BLAST-search of the tags against the IWGSC RefSeq v2.0; the most significant alignments, with high alignment lengths and similarity (>95%), were used to determine physical positions. Out of the three major matches corresponding to three sub-genomes of wheat, the physical position of each SNP was determined based on its genetic map position and the start sequence position (in megabases) from the BLAST alignments. This constructed physical map was then used for localization of identified QTL (Kumar *et al.,* 2024).

### QTL analysis

For the QTL analysis, BLUE values for six spike traits across nine pooled environments (Table 1) were used alongside SBG-SNP genotypic data and the genetic map. Composite interval mapping (CIM; Zeng, 1994) was performed using QTL Cartographer v2.5 (Basten *et al.,* 2002; Wang *et al.,* 2007) to precisely identify QTL locations by accounting for background noise, especially for traits controlled by multiple loci. Cofactors for CIM were selected using forward selection, backward elimination, and stepwise regression (model 6). The genetic contribution (R²) was estimated as the percentage of phenotypic variance explained (PVE). Significant LOD thresholds were set for each trait based on a permutation test with 1,000 iterations (Churchill and Doerge, 1994). QTL were considered significant if they exceeded the LOD threshold in any environment, and the same QTL was detected below the threshold value, but had a LOD score above 2.5 in other environments. QTL identified in at least 50% of the environments were labeled as stable. These QTL were mapped onto the physical map using MapChart 2.2 (Voorrips, 2002).

### Candidate genes identification and *in-silico* expression analysis

Candidate genes (CGs) associated with each QTL were identified within a 1.0 Mb region upstream and downstream of the QTL’s peak. A 1.0 Mb window around the QTL peak is a commonly accepted criterion (Kumar *et al.,* 2021; Pal *et al.,* 2022; Singh *et al.,* 2023) to capture relevant genomic regions, increasing the likelihood of identifying functionally relevant genes (Edwards and Batley, 2010). CGs were identified using BioMart in the EnsemblPlants database for *Triticum aestivum* (https://plants.ensembl.org/biomart/martview). *In silico* gene expression analysis was carried out using four separate transcriptome datasets from the WheatExpression Browser, powered by WheatOmics (Ma *et al.,* 2021; http://wheatomics.sdau.edu.cn/).

Although some datasets included RNA-seq data from multiple abiotic stresses, only heat stress data was used for in-silico expression analysis. The datasets used were: (a) heat and drought stress dataset from Liu *et al.,* (2015), with RNA-seq data from wheat cv. TAM107 seedlings exposed to heat stress (40°C); (b) RNA-seq data from leaf, root, and grain tissues under heat stress during grain filling, from two wheat cultivars, Atay85 and Zubkov (https://www.ncbi.nlm.nih.gov/bioproject/?term=PRJNA358808); (c) heat stress dataset for tolerant (HD2985) and susceptible (HD2329) wheat cultivars (Kumar *et al.,* 2015); and (d) heat stress transcriptomes dataset (https://www.ncbi.nlm.nih.gov/bioproject/PRJNA427246). Genes with a fold change (FC) of +2 or −2 TPM compared to the control were considered differentially expressed. Heat maps of gene expression patterns were generated using TBtools (https://bio.tools/tbtools; Chen et al. 2020).

### Identification of heat stress-responsive genes

Candidate genes involved in heat stress tolerance were identified through two approaches: a literature review to find previously reported heat stress-tolerance proteins and analysis of gene networks using the KnetMiner database (http://knetminer.com/Triticum_aestivum/).

### Comparison of identified QTL with reported meta-QTL

The QTL identified in this study were compared to meta-QTL (M-QTL) from four previous heat stress-related studies (Acuña Galindo *et al.,* 2015; Liu *et al.,* 2020; Kumar *et al.,* 2021; Yang *et al.,* 2024) by matching the physical positions of associated markers with the coordinates from CerealDB (https://www.cerealsdb.uk.net/) and the JBrowse wheat genome browser (https://wheat-urgi.versailles.inra.fr/Tools/Jbrowse). If the QTL markers overlapped with the M-QTL region, the QTL was considered co-localized with the M-QTL.

### Co-localization of QTL and heat-responsive genes

Previously known heat stress-responsive genes were retrieved from WheatOmics (http://wheatomics.sdau.edu.cn/; Ma *et al.,* 2021) and relevant literature. Their physical positions were compared with the QTL regions identified in this study to reveal their co-location with heat-responsive genes.

### Development and validation of KASP marker

To develop Kompetitive Allele Specific PCR (KASP) marker for individual SNP markers (Supplementary Table S2), two locus-specific primers and a common primer were generated using PolyMarker (http://www.polymarker.info/). The KASP assay involving two locus-specific primers attaching the standard FAM and HEX tails at 5’ end and the target SNP at the 3’ end, and a common primer were together synthesized by Integrated DNA Technologies (IDT, India). Laboratory protocols for the preparation of KASP assay mix and PCR conditions were used as per the recommendation of LGC Genomics (http://www.lgcgroup.com/). The KASP genotyping assay was tested/validated on a subset of DH lines (segregating for the corresponding marker) in 96-well plate having a final reaction volume of 10 μl containing of 2 μl (25 ng/μl) genomic DNA, 5.0 μl KASP mix, 0.14 μl of KASP primer mix, and 2.86 μl ddH_2_O. On each SNP reaction plate, at least two sample with no-template were included as control. The PCR reactions were performed in BioRad CFX-96 RT-PCR thermal cycler with the following profile: initial hot-start step 94 ^0^C for 15 min, followed by 10 touchdown cycles (94 ^0^C for 20s, touchdown at 61 ^0^C to 55 ^0^C, decreasing 0.6 ^0^C per cycle, 60 s) and 26 cycles of amplification at 94 ^0^C for 20 s and 55 ^0^C for 1 min. Allele discrimination based on relative fluorescence unit (RFU) data of amplified products were detected and analyzed using CFX Connect Real-Time PCR detection system (Bio-Rad^®^ Laboratories Inc.).

### Pangenome-assisted development of functional

#### markers for selected candidate genes

The coding DNA sequences (CDS) of the eight important candidate genes having a role in heat stress tolerance were retrieved in FASTA format from Ensembl Plants. Each CDS was used as a query for BLASTn against the whole genome sequences of each of the 17 wheat cultivars (including the Chinese Spring) used earlier to develop the wheat pangenome (https://plants.ensembl.org/Triticum_aestivum/Info/Strains?db=core). The resulting aligned sequences for each gene across the cultivars were subjected to multiple sequence alignment using MultAlin (http://multalin.toulouse.inra.fr/multalin/) to identify structural variations such as SNPs and InDels. Further, SSRs were identified in each candidate gene using the MISA tool (https://webblast.ipk-gatersleben.de/misa/). Gene structures were visualized by uploading CDS and corresponding genomic sequences to the Gene Structure Display Server (GSDS; https://gsds.gao-lab.org/).

### Primer design and PCR analysis of SSRs

Details of primer sequences of SSRs used in the present study are given in Supplementary Table S3. Candidate gene sequences were used to design gene-specific microsatellite (SSR) primers (forward and reverse) using Primer3Plus tool (https://www.primer3plus.com/). Appropriate primer sequences were selected and synthesized (Integrated DNA Technologies (IDT, India) for their utilization in targeted amplification of the genomic region during molecular profiling.

The genomic DNA extraction of a set 16 wheat genotypes including the two parents of the DH population used during the present study (Supplementary Table S4) and that of the DH mapping population was carried out essentially in the same manner as described in (Kumar *et al.,* 2024). PCR was performed in a 10 µl reaction mixture containing 10 ng of template DNA, 5µl master mix (Servicebio, BioServe Biotechnologies, Hyderabad), 1 µl each of 10 mM forward and reverse primers and 3µl double-distilled water to make up the volume to 10 μl, using a 96-well Thermal Cycler (Applied Biosystems, Thermo Fischer Scientific, USA). PCR profile was as follows: initial denaturation for 3 min at 94°C (1cycle), denaturation at 94°C for 30 s, annealing at 58°C for 45 s (30 cycle), extension 72°C for 1 min, final extension at 72°C for 10 min at the end hold was at 10°C. The PCR product were run on 2.0% agarose gel at 100 V (250 amp) for 45 min on a electrophoresis system (MS) and the amplified products were visualized under UV light in a gel-doc (Nu-Genius XE, Syngene).

## Results

### Temperature and terminal heat stress during field evaluation

Figure 1 illustrates the daily minimum (Tmin, °C) and maximum (Tmax, °C) temperatures from heading to physiological maturity under the three sowing conditions (TS, LS, and VLS) at Meerut over the 2017–2020 crop seasons. In the reproductive and maturity phases (from heading to physiological maturity), temperatures under the TS condition were optimal with Tmin averaged between 11.6°C and 13.6°C; Tmax ranged from 24.9°C to 28.9°C across the three seasons. Under LS, Tmin averaged between 14.3°C and 16.6°C, and Tmax ranged from 27.9°C to 32.4°C. Under VLS, Tmin ranged from 17.2°C to 18.3°C, while Tmax ranged from 32.4°C to 35.5°C. Additionally, the time from 50% heading to physiological maturity in TS took 62 days, compared to 42 and 38 days in LS and VLS. The duration of the physiological maturity phases at Meerut under TS, LS, and VLS was 152, 121, and 104 days, respectively, with reductions of 31 days (20.4%) under LS and 48 days (31.6%) under VLS.

**Fig. 1.**
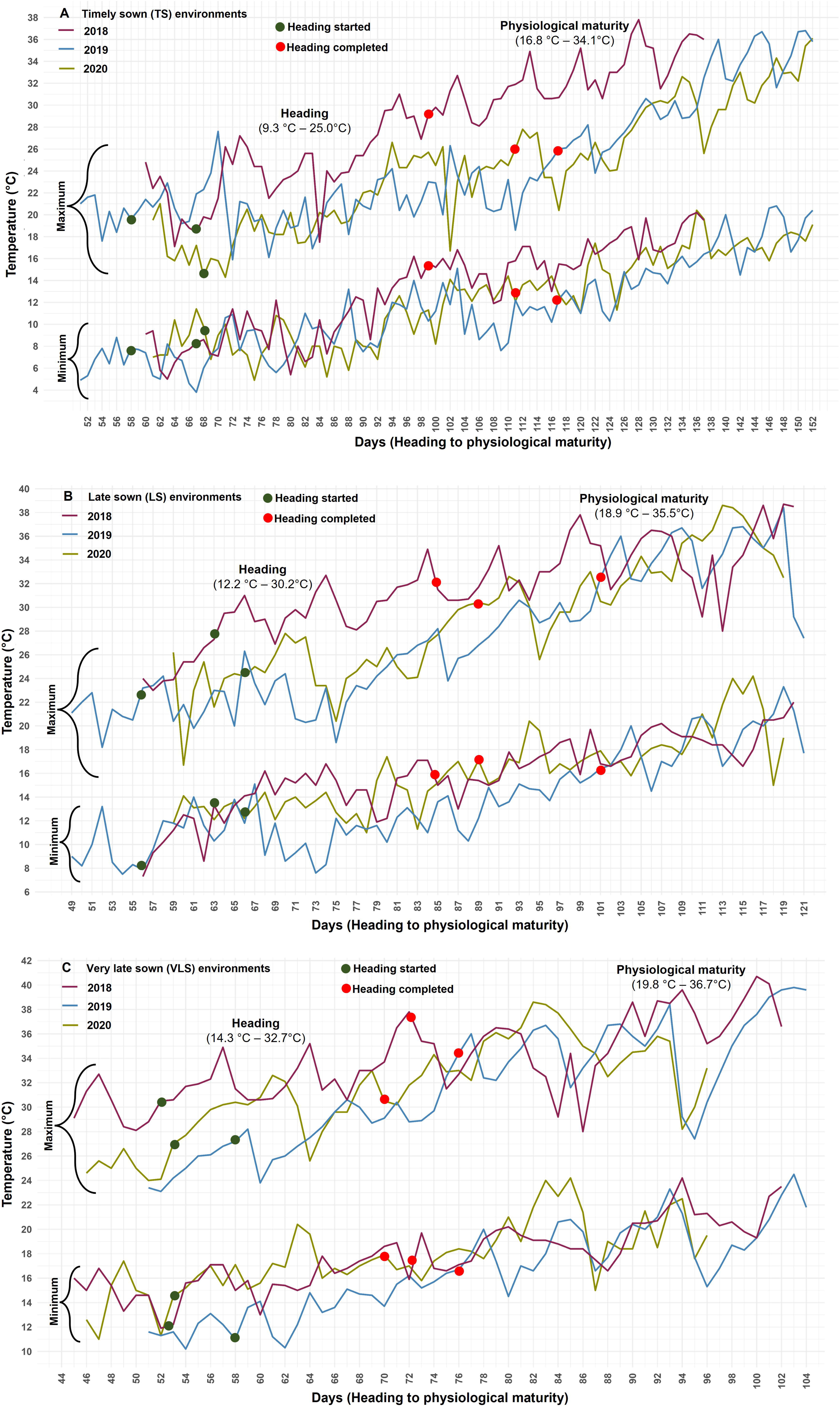
Daily minimum and maximum temperature (°C) fluctuations in (A) timely sown, (B) late sown, and (C) very late sown environments at Meerut location. Y-axis and X-axis respectively represents temperature and number of days after sowing.

At Lucknow, daily Tmin and Tmax were 2–3°C higher across the 2018 and 2019 crop seasons for all sowing conditions (Supplementary Fig. S1). Like Meerut, the temperatures in the TS environment were optimal, with Tmin ranging from 14.3°C to 16.1°C and Tmax from 28.5°C to 31.9°C. Under LS, Tmin ranged from 17.0°C to 19.0°C, and Tmax from 31.7°C to 35.0°C. In VLS, Tmin ranged from 19.7°C to 20.1°C, while Tmax was between 35.5°C and 35.7°C. Overall, the higher Tmin and Tmax during LS and VLS (heat stress environments) at both locations exposed the crop to terminal heat stress, reducing the duration of the reproductive and maturity phases under heat stress conditions (LS and VLS).

### Descriptive statistics (CV%, heritability (_bs_), Ridgeline plots, ANOVA)

The combined BLUE values of the DH lines across the three sowing environments (TS, LS, and VLS) and both locations showed a broad variation for all six spike traits (Table 1). The coefficient of variation (CV%) was low for most traits, except for FF and GWS, which had moderate CV% across all three sowing environments (Supplementary Table S5). Nevertheless, broad-sense heritability for the different traits ranged from moderate to high in all environments.

Ridgeline plots of the DH lines displayed continuous variation and an almost normal distribution, along with transgressive segregation, for all six spike traits (Fig. 2). Both the ridgeline plots and overall mean values indicated a significant decline in phenotypic expression in five traits (except SD) when comparing LS to TS environments, with reductions ranging from [4.70% (NSS) to 18.91% (GWS)]. Similarly, declines were observed in VLS compared to TS environments [4.06% (SD) to 31.57% (GWS)] and in VLS compared to LS environments [1.57% (SD) to 15.62% (GWS)] (Fig. 3). In contrast, SD exhibited a positive change under both LS and VLS conditions compared to TS, likely due to the disproportionate decline in SL and NSS under the two heat stress environments. The combined ANOVA showed significant differences among genotypes, sowing environments, locations, years, two-way, three-way and four-way interactions for all the six spike traits with rare exceptions (Supplementary Table S6).

**Fig. 2.**
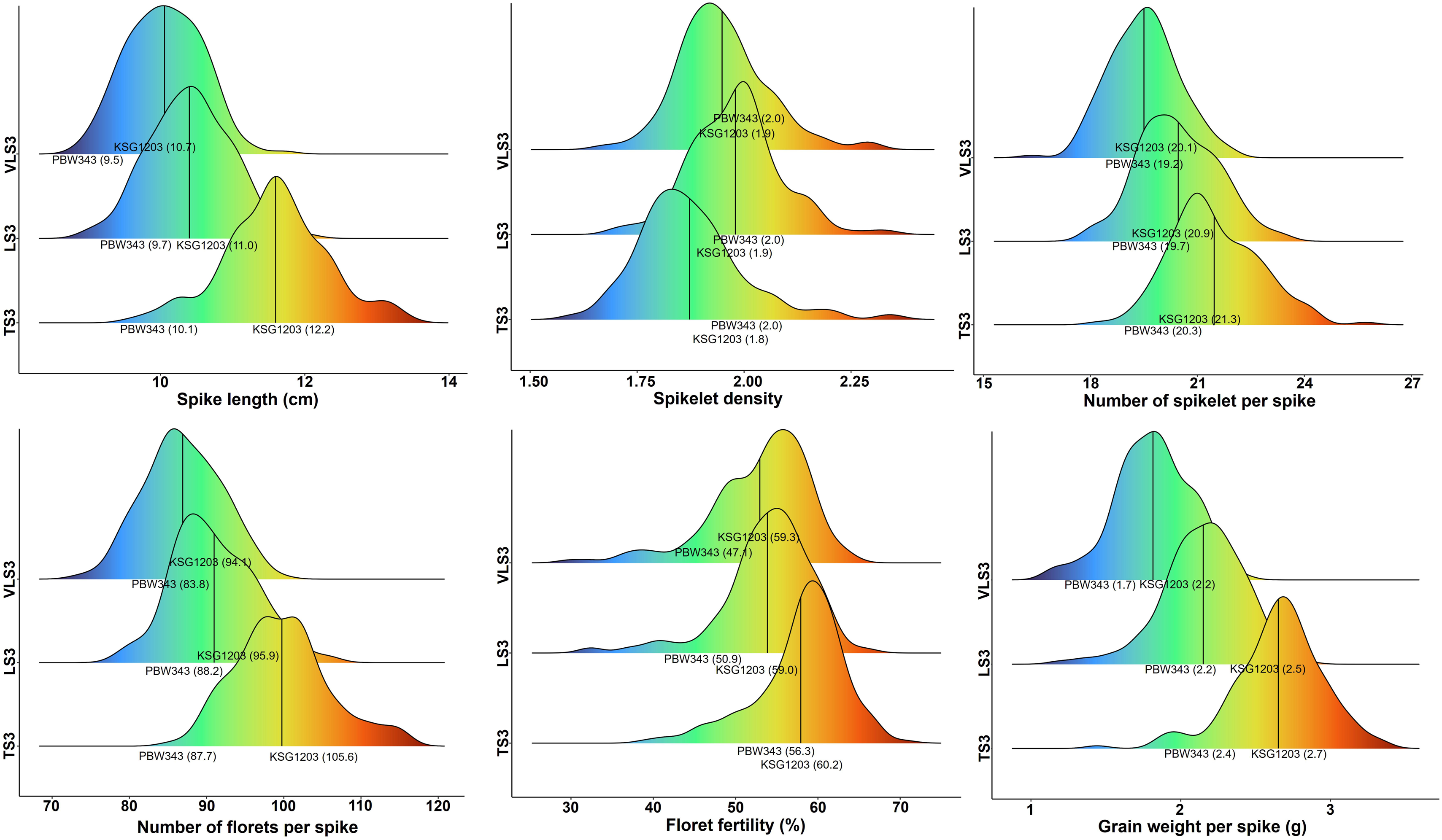
Ridgeline plots drawn using overall pooled data of three sowing environments (TS3, LS3, and VLS3; for details refer Table 1). Each of the three plots depict distribution of six spike related traits in the doubled-haploid wheat mapping population evaluated under three sowing environments (TS: timely sown, LS: late sown, VLS; very late sown).

**Fig. 3.**
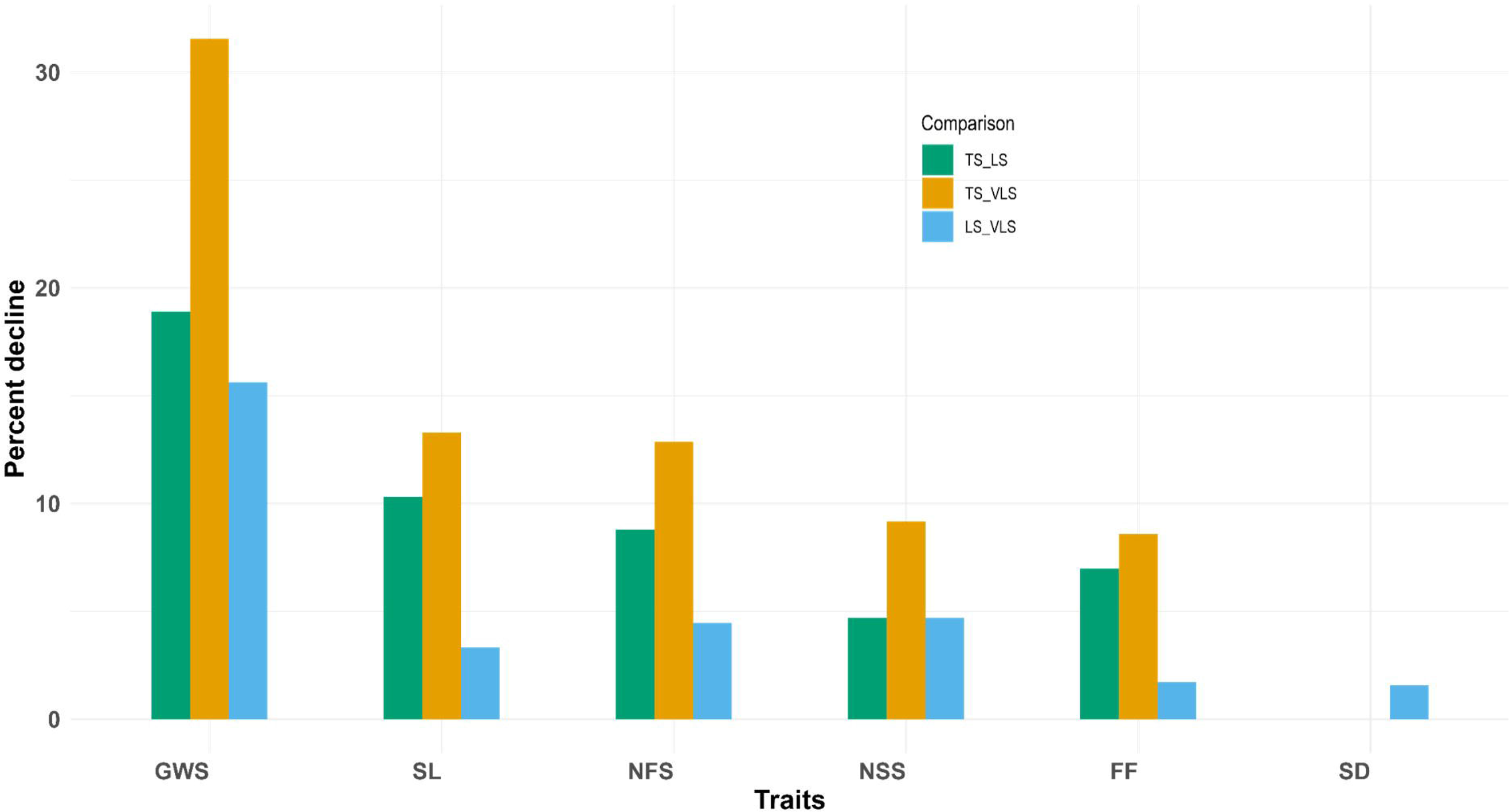
Percent (%) decline in the mean values of DH lines for the six spike traits under late sown (LS) and very late sown (VLS) environments in comparison to the timely sown (TS) environment.

### Correlations and path-coefficient analysis

The correlation coefficients for the 15 trait pairs across each of the TS, LS, and VLS environments were calculated using pooled BLUE values over three years and two locations (Supplementary Fig. S2). In the LS and VLS environments, eight trait pairs had significant positive correlations, while in the TS environment, six pairs exhibited positive correlations. On the other hand, four pairs in LS and VLS, and five pairs in TS showed significant negative correlations, with the rest being non-significant. Particularly, FF and GWS, as well as NFS and NSS, consistently showed strong positive correlations across all three sowing conditions.

The path coefficient analysis showed consistent patterns of direct contributions from five independent traits to GWS across both TS and LS environments. Traits such as NSS, NFS, and FF had positive direct contributions, while the remaining two traits showed negative direct effects. In both environments, FF had the highest positive contribution, followed by NFS and NSS (Supplementary Table S7). Additionally, SL indirectly contributed to GWS through SD, NFS, NSS (LS only) in both environments. In LS, SL, SD and NFS also made notable positive indirect contributions to GWS via NSS. In both environments, all traits except FF made positive indirect contributions to GWS through NFS, with SL and NSS being the highest contributors. In the VLS environment, four of the five traits (excluding NSS) had positive direct effects on GWS, with FF, NFS, and SL contributing the most. Apart from FF, the remaining traits made positive indirect contributions to GWS via NFS. However, the indirect contributions of certain traits were either small or negative.

### Multi-environment QTL mapping

The findings from the QTL analysis of six spike traits across nine different environments (three sowing conditions: TS, LS, and VLS, pooled separately for Meerut and Lucknow, as well as pooled across both locations; Table 1) are outlined in Table 2. A total of 51 QTL were identified, spanning 19 chromosomes (including ChrUn), except for chromosomes 1D, 4B, and 4D. The QTL, with LOD scores ranging from 3.4 to 7.6, explained between 7.1% and 23.6% of the phenotypic variation. NSS and FF had the fewest QTL (7 each), while GWS had the most (11). Five QTL each were located on chromosomes 3A, 4A, 5A, 5B, and 6B, while individual QTL were identified on chromosomes 1A, 3D, 6A, 6D, and 7D. Some QTL were co-located including two each on chromosomes 2A, 2D, 3A, 4A, 5A, 5B, and 5D, and two pairs on chromosome 6B (Fig. 4a; Supplementary Fig. S3). Many QTL were specific to particular sowing environments (TS=17, LS=10, VLS=18), with six QTL shared between environments. Out of the total 51 QTL, two QTL, one each for SD (*QSd.ccsu-5D*) and NSS (*QNss.ccsu-3A.2*) were stable in TS environment whereas eight QTL in LS (*QSl.ccsu.4A*, *QNss.ccsu-6B.2*, *QSd.ccsu-6B*, *QGws.ccsu-4A*, *QGws.ccsu-5B.2*, *QSl.ccsu-5D*, *QNss.ccsu-6B.1*, and *QFf.ccsu-5B*) and four QTL in VLS (*QNfs.ccsu-2D*, *QSl.ccsu-6A*, *QSd.ccsu-4A*, and *QFf.ccsu-3A*) environments were stable. The favourable alleles were contributed by KSG1203 for 21 QTL and by PBW343 for the remaining 30 QTL.

**Fig. 4.**
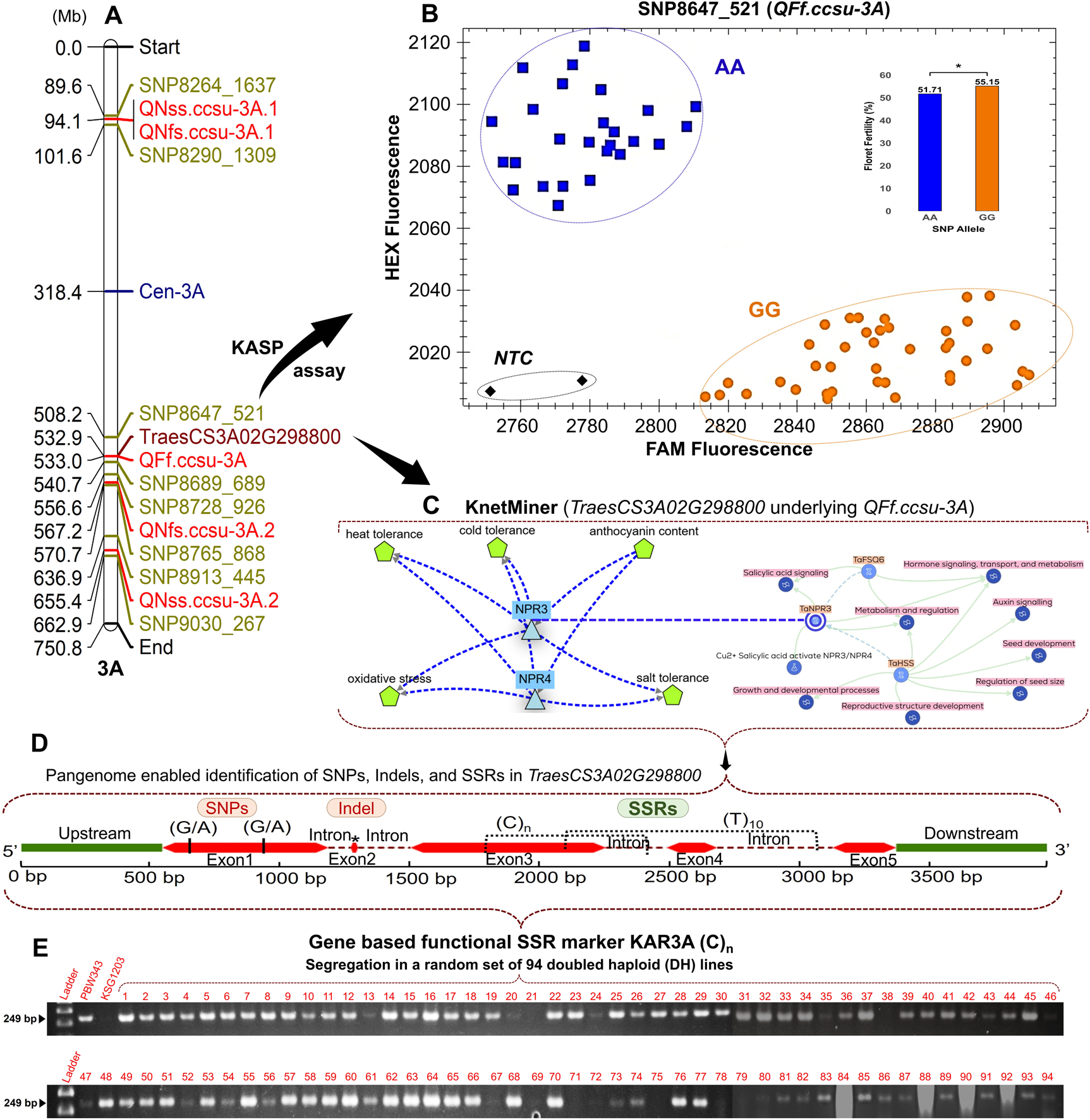
Integrated analysis of QTL *QFf.ccsu-3A* for floret fertility and its underlying candidate gene TraesCS3A02G298800 on chromosome 3A. (A) Physical position of QTL-associated SNPs and corresponding candidate gene; (B) KASP assay for SNP8647_521 discriminating clear genotyping clusters (AA and GG) and phenotypic effect on floret fertility; (C) KnetMiner network analysis linking TraesCS3A02G298800 through NPR3/NPR4 gene with heat stress tolerance traits; (D) Gene structure showing identified SNPs, Indels, and SSRs in the TraesCS3A02G298800 region; (E) Validation of a gene-based functional SSR marker KAR3A (product size 249 base pair) using subset of 94 doubled haploid lines of the PBW343/KSG1203 mapping population.

**Table 2.**
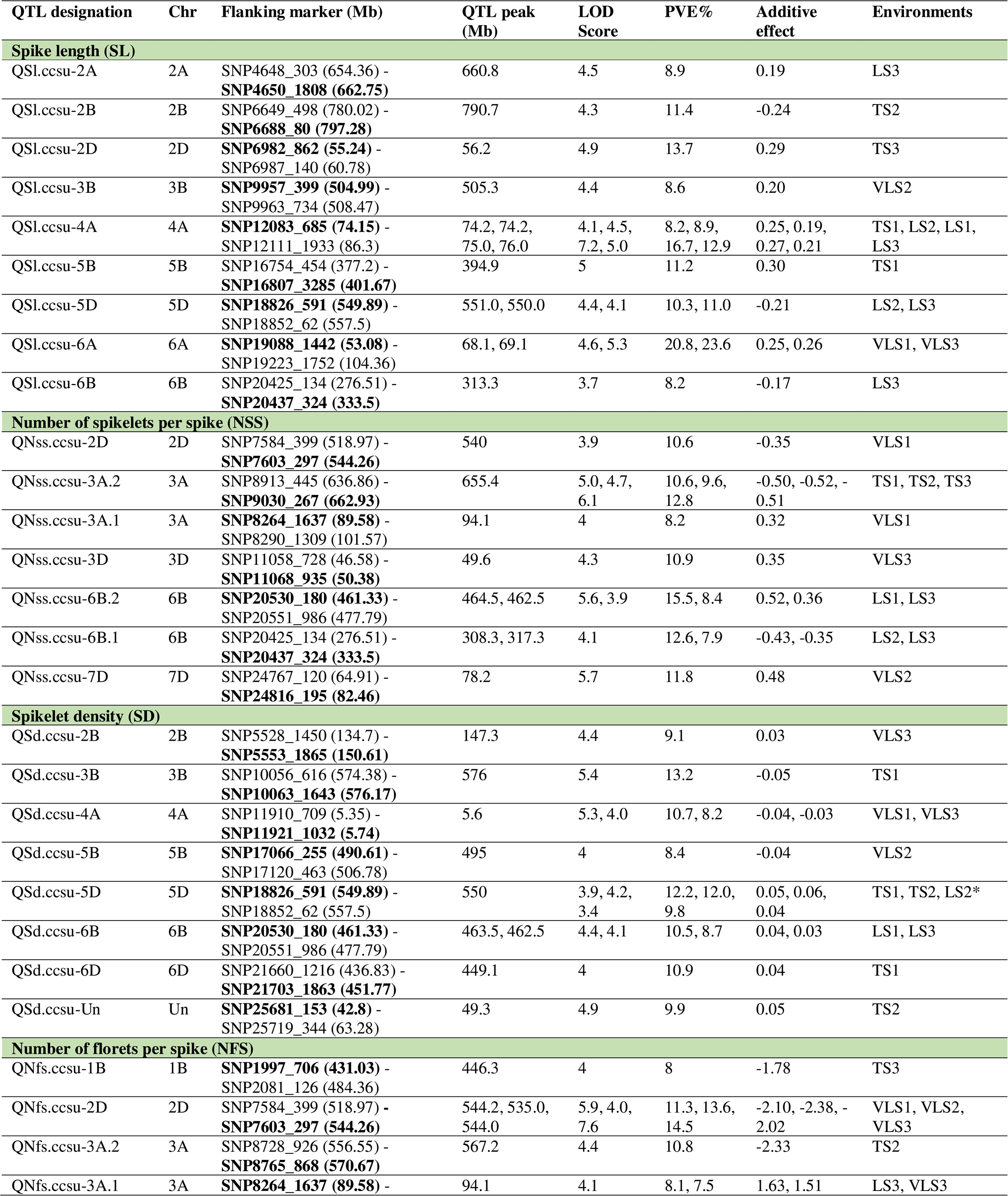
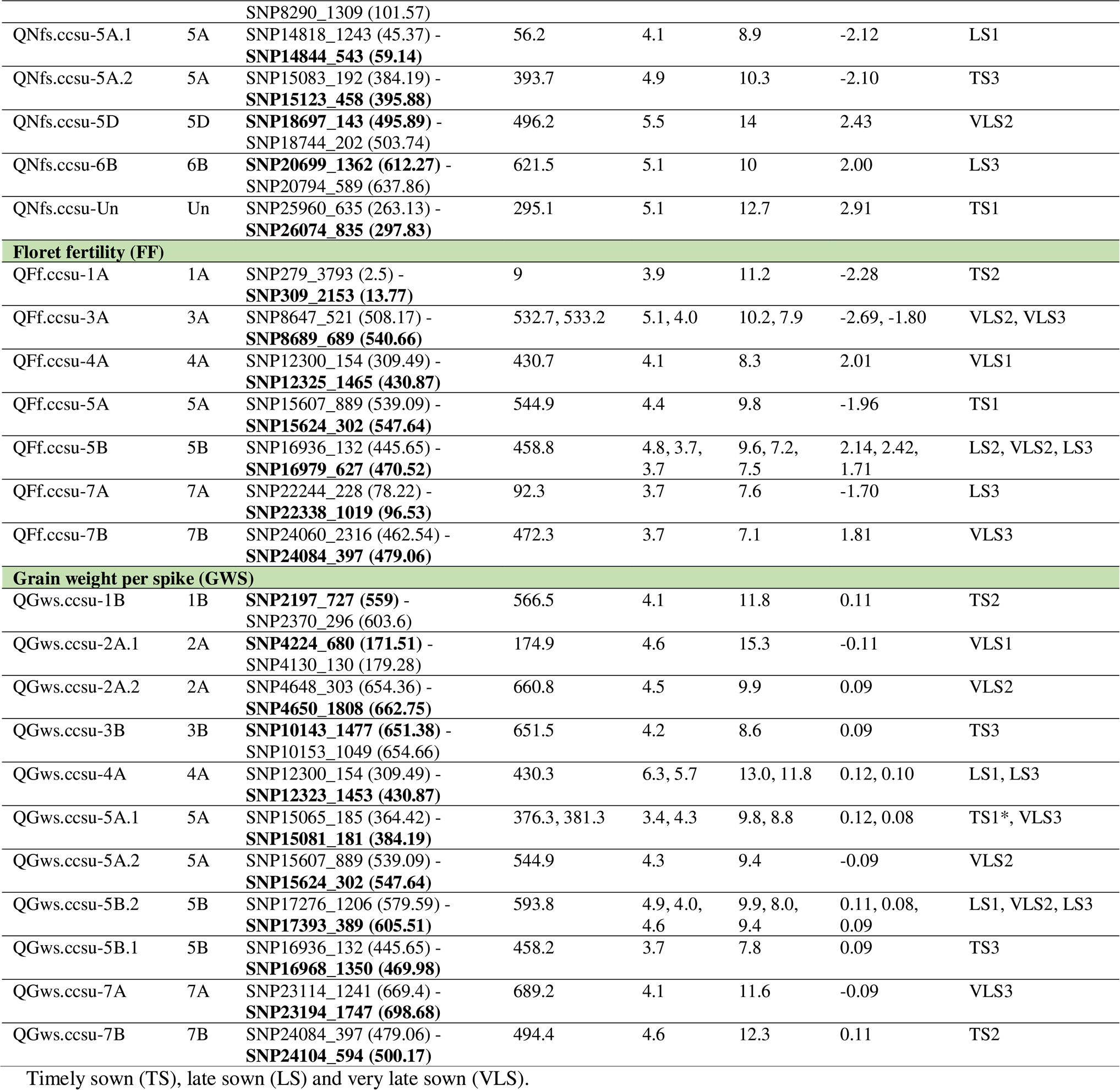
Details of 51 QTL for six spike traits identified in nine pooled environments based on field trials of PBW343/KSG1203 doubled haploid mapping population conducted on different dates of sowing, locations and years.

### Co-localization of QTL with meta-QTL and new QTL

Of the 51 QTL identified for six traits, 42 were co-located with 48 previously reported meta-QTL (Supplementary Table S8), spanning 15 chromosomes (Acuña Galindo et al. 2015; Liu et al. 2020; Kumar et al. 2021; Yang et al. 2024). The remaining nine QTL, reported for the first time in this study, are considered novel. These include three QTL for SD (*QSl.ccsu-2B*, *QSd.ccsu-6D*, *QSd.ccsu-Un*), two for SL (*QSl.ccsu-2B*, *QSl.ccsu-5D*), and one each for NSS (*QNss.ccsu-7D*), NFS (*QNfs.ccsu-Un*), FF (*QFf.ccsu-7B*), and GWS (*QGws.ccsu-7B*). Detected in one to three of the nine pooled environments, these novel QTL had LOD scores between 3.4 and 5.7 and PVE ranging from 7.1% to 12.7%.

### Co-localization of QTL with previously reported genes

This study found QTL that overlapped with six established genes: *TaHsfC2a-B*, *TaCol-B5*, *WAPO1*, *Starch synthase I*, *TaGW2-B1*, and *PI1-1B/WPI-1-1B*. These genes are located within six QTL regions (*QGws.ccsu-7B*, *QGws.ccsu-7A*, *QNss.ccsu-7D*, *QNss.ccsu-6B.1*, *QSl.ccsu-6B*, and *QNfs.ccsu-1B*) distributed on five different chromosomes (1B, 6B, 7A, 7B, and 7D), related to traits such as NFS, NSS, SL, and GWS (Table 3; Supplementary Fig. S3). These genes play roles in heat shock response, starch synthesis, spikelet number per spike, spike architecture (including spikelet and floret development), grain weight, grain yield, and various aspects of spike and grain development (Table 3).

**Table 3.**
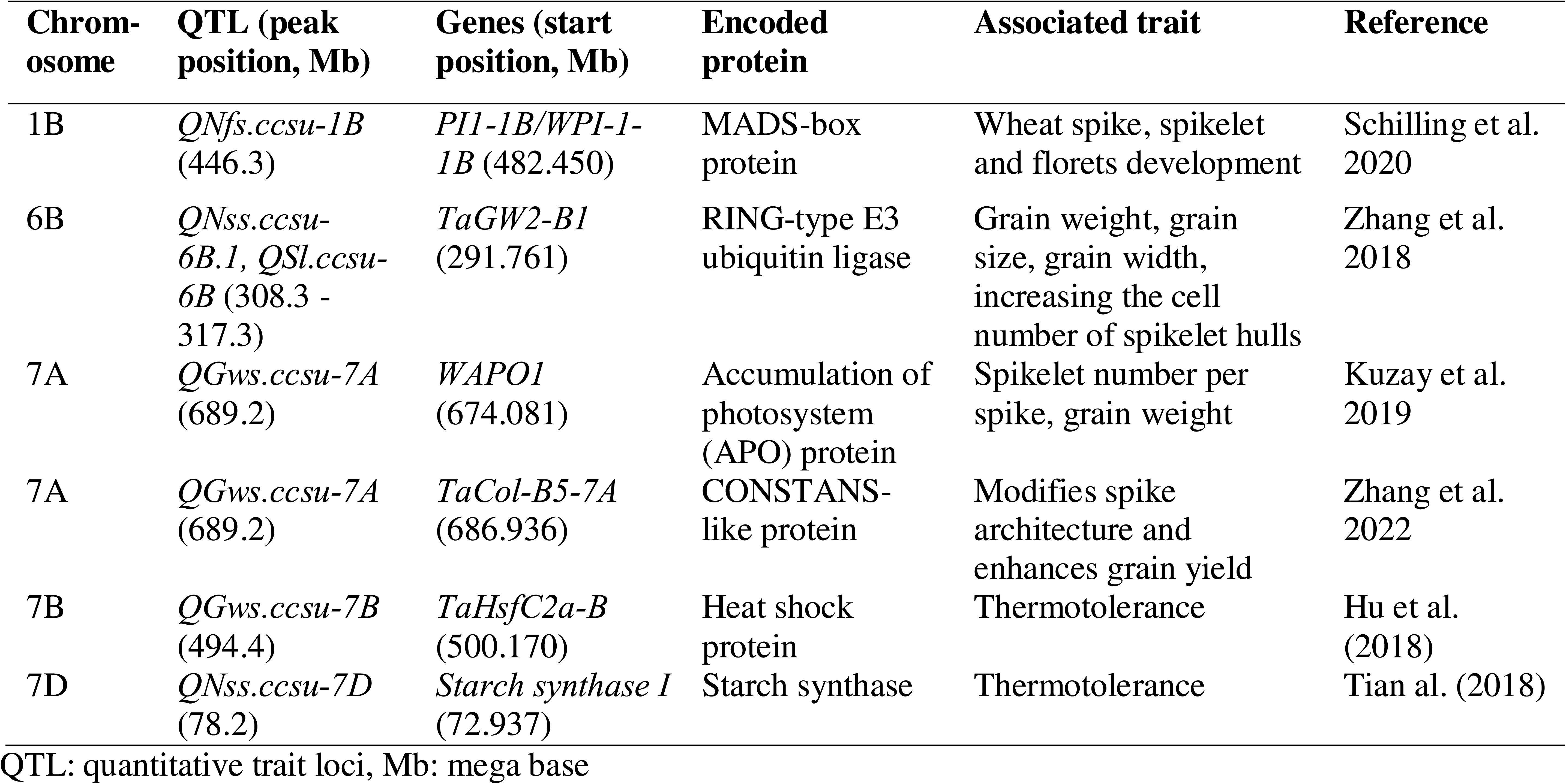
Six known genes and encoded protein for heat tolerance/spike related traits underlying six QTL regions on five different chromosomes of wheat identified during the present study.

### Favourable SNP alleles for selected QTL in three DH lines with highest GWS

To validate the QTL chosen for marker-assisted selection (MAS) (as discussed later), an analysis was conducted to assess the composition of favorable alleles linked to the closest SNP markers associated with the selected QTL in the three DH lines with highest GWS across the TS, LS, and VLS environments. These top-performing DH lines had GWS values (based on pooled data), that surpassed the parent cultivars with highest GWS in each of the respective sowing environments.

In the TS environment, only one QTL related to NSS was chosen for MAS. The favorable allele for the nearest SNP marker linked to this QTL was found in the two of three DH lines with highest GWS, while the other one line did not carry this favourable allele (Table 4). In the LS environment, seven QTL for various spike traits were selected for MAS, and the analysis of the favorable alleles for the corresponding SNP markers showed that the selected lines had four to six favorable alleles. The two DH lines (DH-197 and DH-156) with the highest number (6 each) of favorable alleles had the second and third-highest GWS in the LS environment.

**Table 4.**
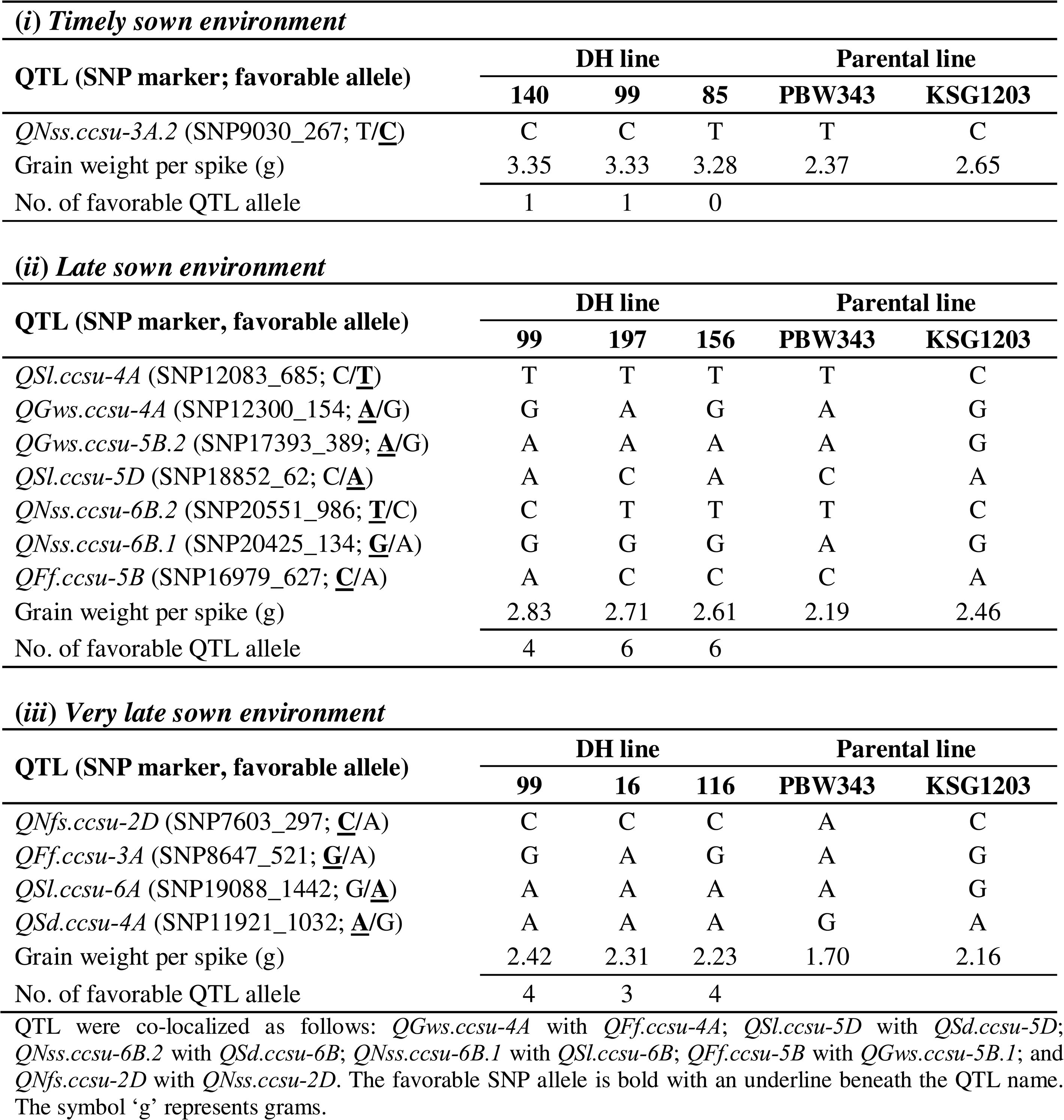
Combinations of favourable SNP alleles in three highest grain weight doubled-haploid (DH) lines of PBW343/KSG1203 population in timely, late and very late sown environments.

A similar analysis was carried out for the top three DH lines in the VLS environment to examine the composition of the favorable alleles associated with the SNP markers flanking the four selected QTL (Table 4). The analysis revealed that the lines contained between three and four favorable alleles, with the top-performing DH line (99) having the highest number of favorable alleles (4), while the other two lines each contained three and four favorable alleles.

### Candidate genes, *in silico* expression, and gene network analyses

We identified 1,183 candidate genes linked to 46 of the 51 QTL across six spike traits analyzed in this study. From this, we selected 465 genes (39.3%) with at least a two-fold expression increase for further examination. A comprehensive review of the literature and the KnetMiner database suggested that 181 of these genes may be involved in abiotic stress responses (Supplementary Table S9), which became the focus of our subsequent analyses. *In silico* expression analysis of these 181 genes revealed fold changes ranging from −64.62 to +15.13 across four distinct expression datasets, including various tissues and developmental stages (see Materials and Methods; Supplementary Table S9). Heat maps of key differentially expressed genes are presented in Supplementary Fig. S4. Data integration from the KnetMiner database and prior literature identified 70 high-confidence genes encoding 33 different proteins involved in complex biological processes, contributing to heat tolerance in plants (Fig. 5). The knowledge network analysis identified eight important candidate genes (Fig. 4c and Supplementary Fig. S5) underlying eight QTL for each of the six spike traits. These genes are involved in providing abiotic stress tolerance including heat tolerance and are involved in controlling traits like grain weight/size/number, spikelet number, spikelet fertility, harvest index, grain yield, etc. Figure 4c shows an example of a KnetMiner knowledge network, highlighting the role of a CG TraesCS3A02G298800 *i.e.* NPR3/NPR4 (532.9 Mb), underpinning a QTL (*QFf.ccsu-*3A; 533 Mb) for FF on long arm of chromosome 3A (3AL).

**Fig. 5.**
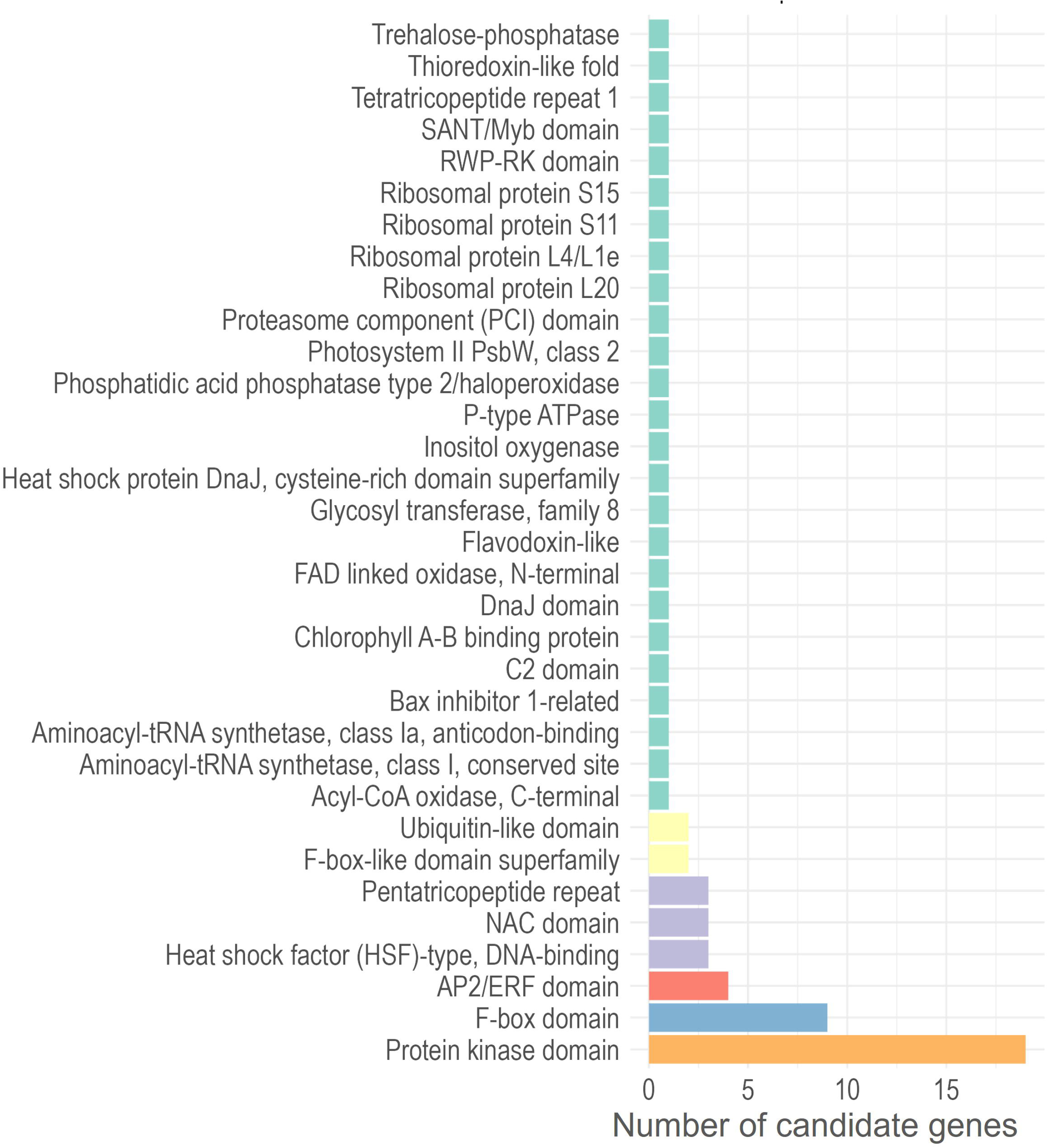
Histogram showing frequencies of candidate genes encoding 33 different proteins involved in heat tolerance.

### KASP markers validation

The SNPs associated with the important QTL could serve as valuable resources for the development of KASP assays. In the present study, efforts were made to develop KASP assays for 12 SNPs, one each associated with 12 important and stable QTL selected for marker-assisted selection (Supplementary Table S2). Among these, KASP assay for a solitary SNP, namely SNP8647_521 associated with a QTL for FF (*QFf.ccsu-3A*) detected in VLS environment showed significant phenotypic effect and segregated in the DH population (Fig. 4b). The favorable allele of the above SNP is responsible for significantly enhancing the FF by 3.44% under the VLS environment. The remaining 11 SNPs, nine were either monomorphic and/or produced indistinguishable clusters, and assays for two SNPs could not be designed.

### Development of gene-based functional

#### markers using pangenome

We were able to identify SNPs, Indels and SSRs in eight important CGs (Supplementary Fig. S6; Supplementary Table S10), using information from 17 wheat cultivars included in pangenome. One or more SNPs were identified in six CGs, a single Indel in seven genes and one to four SSRs in five of the eight genes; these sequence variations occurred upstream, exonic and intronic regions (Fig. 4d; Fig. 6). In the present study, we tried to detect polymorphism due to one SSR each from the five individual genes in a set of 16 wheat genotypes including the two parents of the DH mapping population used during the present study. Five of these SSRs belonging to five individual CGs which include the mononucleotide SSR (C)_n_ (in TraesCS3A02G298800), trinucleotide SSR (GCG)_5_ (in TraesCS4A02G008000), di-nucleotide SSR (CT)_n_ (in TraesCS5D02G445100) and (CT)_7_ (in TraesCS6B02G257400), and a trinucleotide SSR (GCA)_6_ (in TraesCS3A02G298800) were successfully amplified in a set of 16 wheat genotypes; all these five SSRs showed +/− polymorphism (Fig.6 b, d, e, f and h). Further, the SSR KAR3A (C)_n_ (in TraesCS3A02G298800) and (CT)_7_ SSR (in TraesCS6B02G257400) also detected +/− polymorphism between the two parental genotypes (KSG1203 and PBW343) of our DH population (Fig. 6b and f). These two SSRs also segregated in our DH population (Figs. 4e and 7). The segregation of SSR (CT)_7_ showed a good fit to 1:1 segregation (Chi-square _(1:1)_ = 0.17) in the DH population (Fig. 7), as expected, whereas the SSR KAR3A (C)n showed segregation distortion in the DH lines (Chi-square _(1:1)_ = 40. 25).

**Fig. 6.**
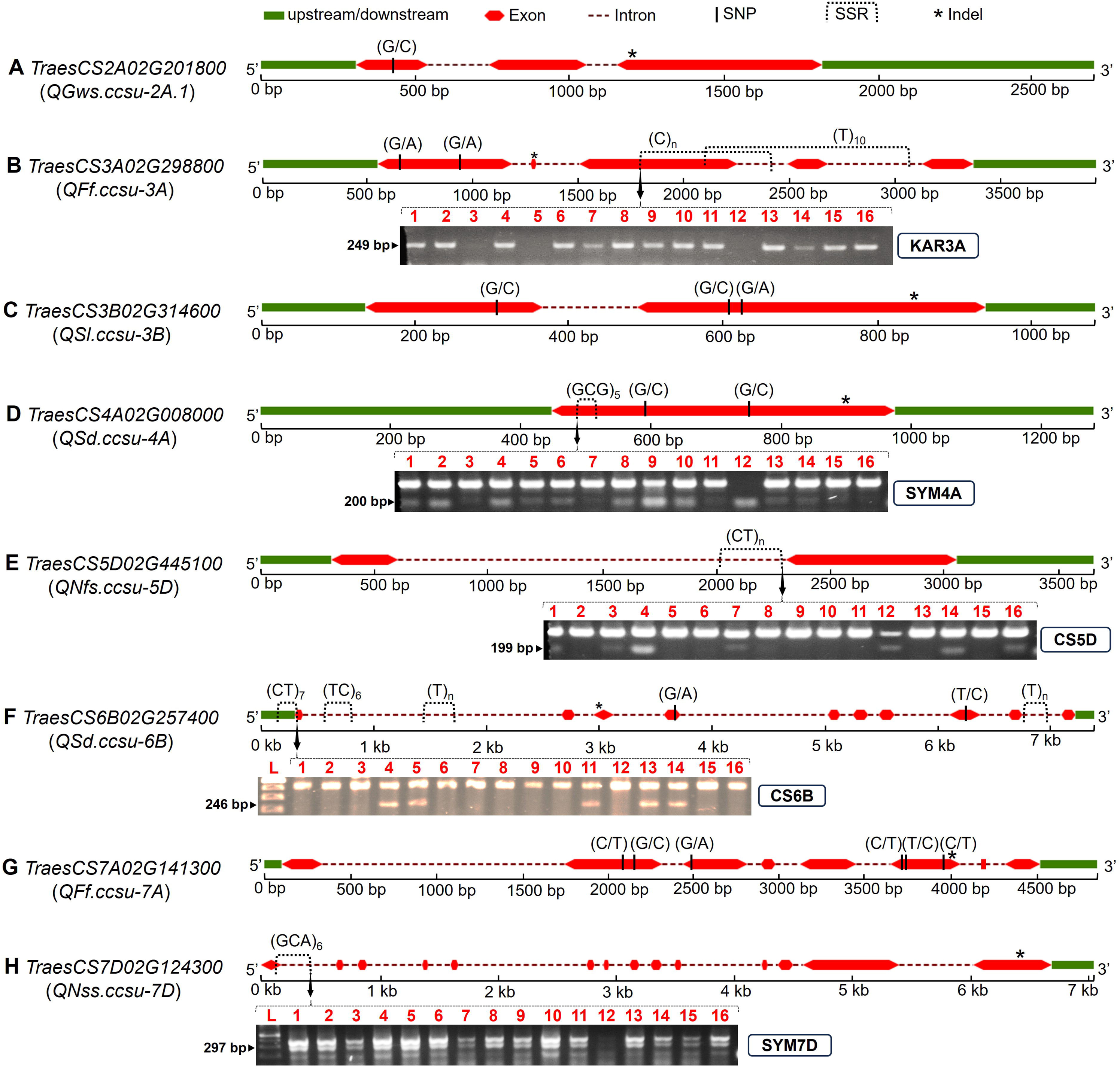
Structures of eight candidate genes (A-H) underlying QTL on different wheat chromosomes, depicting exons (red), introns (dashed lines), upstream/downstream regions (green), with annotated SNPs, indels (*), and SSRs (dotted boxes). Five gene-based functional SSR markers, namely KAR3A (B), SYM4A (D), CS5D (E), CS6B (F), SYM7D (H) were validated across 16 wheat accessions (1-16): KSG0025, KSG0809, KSG1188, KSG1194, KSG1203, PBW343, KSG1186, KSG1172, KSG1189, KSG1202, IC328433, KSG1177, IC533717, KSG1214, IC416215, and IC290080.

**Fig. 7.**
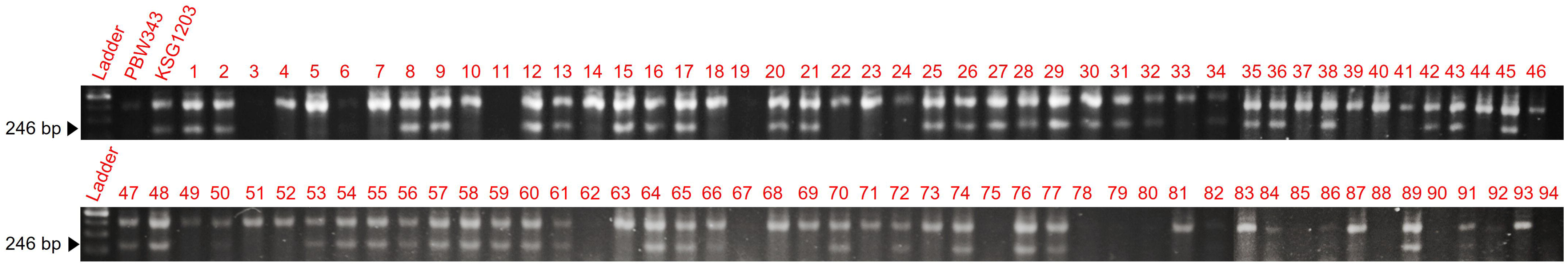
Segregation of SSR CS6B (CT)_7_ (product size 246 base pair) among a random set of 94 (1-94) doubled-haploid lines of PBW343/KSG1203 mapping population.

## Discussion

Climate change has led to increased frequency of both short- and long-duration heat episodes, now affecting nearly half of the global wheat crop. The Indo-Gangetic Plains (IGP) may become unsuitable for wheat cultivation by 2050 due to persistent heat stress, with a 1 °C rise potentially reducing yields by 4–5 Mt (Ortiz *et al.,* 2008; Boraiah *et al.,* 2023). While heat stress affects all growth stages, terminal heat stress during reproductive and maturity phases causes the most severe yield penalties (Parent *et al.,* 2017; Jacott and Boden, 2020; Farhad *et al.,* 2023; Qian *et al.,* 2025). Despite multiple studies on heat tolerance, limited attention has been given to spike-related traits under heat in the IGP (Sharma *et al.,* 2016; Singh *et al.,* 2014; Bhusal *et al.,* 2017; Raveendran *et al.,* 2020; Manjunath *et al.,* 2024). This study, through multi-environment QTL analysis of six spike traits under terminal heat stress, expands previous findings and identifies new QTL, candidate genes (CGs), KASP assay, and gene-based functional markers. These genomic resources offer valuable tools for marker-assisted selection (MAS) to improve grain yield under heat stress conditions in wheat.

### Suitability of mapping population and heat stress environments

The DH population evaluated under timely sown (TS), late sown (LS), and very late sown (VLS) conditions at two locations exhibited significant phenotypic variation, transgressive segregation, and high heritability across six spike traits, indicating valuable QTL contributions from both parents (Table 2; Supplementary Fig. S3). This suggests the potential to combine favorable alleles via selection to improve spike traits and grain yield under heat stress. Mean trait values declined under LS and VLS conditions (Fig. 3), consistent with heat-induced disruptions to spike initiation, reproductive organ development, pollen viability, seed set, and grain filling (Prasad *et al.,* 2008; Farooq *et al.,* 2011; Sehgal *et al.,* 2018; Bheemanahalli *et al.,* 2019; Abdelrahman *et al.,* 2020a; Jacott and Boden, 2020; Jagadish, 2020; Ullah *et al.,* 2024; Qian *et al.,* 2025), contributing to yield reductions as reported previously (Kumar *et al.,* 2024).

Grouping environments by sowing time (TS, LS, VLS) aligned with temperature trends during heading to maturity, validating the pooling strategy. Grain weight per spike (GWS) declined by 18.91% in LS vs. TS, 31.57% in VLS vs. TS, and 15.62% in VLS vs. LS— consistent with yield reductions of 25–60% reported in earlier studies (Gupta *et al.,* 2012; Bal *et al.,* 2022; Sareen *et al.,* 2023; Li *et al.,* 2023). Collectively, these results confirm that the chosen population and stress environments were appropriate for dissecting the genetic control of spike traits under terminal heat stress.

### QTL detection and comparison with previous studies

A total of 51 QTL (LOD 3.4–7.6; PVE 7.1–23.6%) associated with six spike traits were identified across nine pooled environments (Table 1), with the number increasing to 60 when including nine GNS QTL reported previously using the same DH population (Kumar *et al.,* 2024). These QTL were distributed across 19 chromosomes (including ChrUn), excluding 1D, 4B, and 4D, and were integrated into the wheat physical map (IWGSC RefSeq v2.0), allowing precise localization. Physical mapping enabled accurate positioning of QTL, improved cross-population comparisons, and facilitated candidate gene discovery with higher resolution and confidence. This integration enhances the utility of QTL for marker-assisted selection and functional validation in wheat breeding programs.

Previous studies reported QTL for the six spike traits across all wheat chromosomes, including 1D, 4B, and 4D—absent in the current study (Singh *et al.,* 2022; Erena *et al.,* 2021; Telfer *et al.,* 2021). In contrast, prior IGP-based studies focusing on SL and GWS detected QTL on only nine chromosomes (2A, 4B, 4D, 5A, 5B, 6B, 6D, 7B, 7D) (Bhusal *et al.,* 2017; Raveendran *et al.,* 2020; Manjunath *et al.,* 2024). A recent integrative GWAS and meta-QTL study identified 170 loci for SL, SNS, and GNS, with 169 hub genes within 76 of these QTL revealed through transcriptomics, chromatin accessibility, histone modifications, eQTL, and protein interaction analyses—underscoring the complexity of spike trait regulation (Ai *et al.,* 2024). Several spike-related QTL were co-located, suggesting either tight linkage or pleiotropy, as previously described (Mason *et al.,* 2011). Distinguishing pleiotropic from closely linked QTL will benefit from high-resolution mapping and the use of pleiotropy-aware computational models (Knight *et al.,* 2001; Dwivedi *et al.,* 2024).

Among the 51 QTL identified, 17, 10, and 18 were specific to TS, LS, and VLS environments, respectively, while 2, 1, and 3 QTL were shared between TS–LS, TS–VLS, and LS–VLS, respectively—highlighting their relevance for breeding heat-tolerant wheat. Approximately 25% of QTL were stable across pooled environments and consistently associated with the same favorable alleles. The instability of the remaining QTL likely reflects genotype × environment interactions involving sowing time, location, and year effects (Tardieu *et al.,* 2012), though these may still hold value under specific heat stress scenarios.

Pooled environment analysis enhanced the precision and robustness of QTL detection, enabling the identification of both stable and environment-specific QTL. A dual selection strategy was applied to prioritize breeding-relevant QTL: (i) a relaxed LOD threshold of 2.5 captured minor yet potentially valuable QTL, and (ii) QTL detected in ≥50% of environments were classified as stable, ensuring moderate to high environmental consistency (Holland, 2007). QTL meeting these criteria with high PVE are strong candidates for breeding heat-tolerant wheat.

### Trait associations and key environment specific QTL

Correlation and path coefficient analyses indicated that NSS, NFS, FF, and SL positively contributed to GWS under TS conditions. Among the 16 QTL identified for these traits and GWS, only *QNss.ccsu-3A.2* (PVE 9.6–12.8%) was both stable and co-located with a grain yield QTL (*QGy.ccsu-3A*) (Kumar *et al.,* 2024). This QTL also overlapped with a previously reported MQTL (Kumar *et al.,* 2021; Yang *et al.,* 2024), underscoring its robustness and potential utility in MAS to improve GWS under optimal conditions.

Trait contributions to GWS under LS closely mirrored TS conditions, with NSS, NFS, FF, and SL showing positive effects. Of the 14 QTL identified for these traits and GWS in LS, seven were stable and major (high PVE%): *QFf.ccsu-5B*, *QGws.ccsu-4A*, *QGws.ccsu-5B.2*, *QNss.ccsu-6B.1*, *QNss.ccsu-6B.2*, *QSl.ccsu-4A*, and *QSl.ccsu-5D*. Five of these overlapped with unstable QTL, and six were co-located with reported MQTL, confirming their robustness (Kumar *et al.,* 2021; Yang *et al.,* 2024; Tan *et al.,* 2024). *QSl.ccsu-5D*, associated with SL, is novel. Notably, *QGws.ccsu-4A* and *QGws.ccsu-5B.2* co-localized with *QDth.ccsu-4A* and *QTgw.ccsu-5B.2*, respectively, previously reported in the same population (Kumar *et al.,* 2024) highlighting their relevance for improving GWS and overall grain yield.

Under VLS environment, NFS, FF, SD, and SL positively contributed to GWS, while NSS—important in TS and LS—was no longer influential, suggesting a shift in GWS formation mechanisms under severe heat stress. This may reflect changes in source–sink dynamics, enzymatic activity, hormone balance, and metabolite profiles triggered by high temperature (Sehgal *et al.,* 2018; Abdelrahman *et al.,* 2020a, b; Janni *et al.,* 2020; Lal *et al.,* 2021; Shokat *et al.,* 2023; Khanzada *et al.,* 2025; Duan *et al.,* 2025). Of 18 QTL detected for GWS and associated traits, four were stable and major (high PVE%): *QFf.ccsu-3A*, *QNfs.ccsu-2D*, *QSd.ccsu-4A*, and *QSL.ccsu-6A*. QTL *QNfs.ccsu-2D* co-located with *QNss.ccsu-2D*, and *QFf.ccsu-3A* overlapped with *QNgs.ccsu-3A* reported previously using the same population (Kumar *et al.,* 2024). These stable QTL across TS, LS, and VLS, together with those from our prior studies, represent valuable targets for marker-assisted recurrent selection (MARS) to improve grain yield under terminal heat stress.

### Validation of QTL in the DH lines with high GWS

Validation of selected QTL for MARS was performed by examining allele composition in the three DH lines with the highest GWS across TS, LS, and VLS environments (Table 4). In TS, a favorable allele for the single selected QTL was present in two of the top lines. In LS and VLS, these lines carried favorable alleles for 4–6 and 3–4 QTL-associated markers, respectively. As the DH population was not pre-selected for GWS via MAS, not all favorable alleles were expected to occur in high-yielding lines. In contrast, low-GWS lines in each environment carried only 1–3 favorable alleles, supporting the relevance of the selected QTL for GWS improvement.

Some high GWS lines with fewer favorable alleles, such as DH-140 and DH-99, likely benefitted from additional genomic regions. DH-99, the top-performing line in TS and LS, also had the highest GWS under LS and VLS and showed superior performance for SL, FF, NGS, TGW, GFD, BY, and HI across all environments. These findings confirm that lines combining favorable QTL alleles can be developed via MARS to produce early-maturing, high-yielding genotypes adapted to both optimal and heat-stressed conditions (Kumar *et al.,* 2024).

### Candidate/known genes and functional insights

Candidate genes (CGs) were identified within 2 Mb of QTL peaks, a validated approach for localizing functionally relevant genes (Kumar *et al.,* 2021; Pal *et al.,* 2022; Singh *et al.,* 2023). Among 181 CGs linked to abiotic stress tolerance, 70 genes encoding 33 proteins were associated with heat stress tolerance, showing *in silico* expression fold change ranging from −9.01 to +10.79. These included six transcription factor families: F-box, NAC, RWP-RK, AP2/ERF, HSF-type, and SANT/Myb (Fig. 5; Supplementary Table S9).

KnetMiner analysis identified a CG TraesCS3A02G298800, corresponding to NPR3/NPR4 (salicylic acid receptors), underpinning a floret fertility QTL (*QFf.ccsu-3A*) on chromosome arm 3AL. This CG is associated with tolerance to heat, oxidative, cold, and salt stresses, and influences anthocyanin content. The NPR3 also contributes to growth, reproductive structure development, seed size regulation, and salicylic acid/auxin signaling. NPR3 and NPR4 act as adaptors for CUL3-based E3 ligases, which mediate NPR1 degradation in the presence of salicylic acid, thereby negatively regulating SA signaling (Yan and Dong, 2014). This regulation is crucial for modulating SA responses under biotic and combined stresses, indicating their potential role in abiotic stress tolerance. (Ban and Estelle, 2021).

Additionally, CG TraesCS7D02G124300, associated with *QNss.ccsu-7D* on 7DS, interacting with *TaEYE* (oxidative stress), *TaSPI* (grain and spikelet number, HI, canopy temperature), and *OsSIZ1* (heat/drought tolerance, spikelet fertility and seed length) (Supplementary Fig. S5). Remaining six CGs, each linked to distinct QTL (*QFf.ccsu-7A*, *QGws.ccsu-2A*.*1*, *QNfs.ccsu-5D*, *QSd.ccsu-4A*, *QSd.ccsu-6B,* and *QSl.ccsu-3B*), were implicated in heat and oxidative stress responses (Supplementary Fig. S5; Supplementary Table S9). Three of these, namely TraesCS2A02G201800 (*QGws.ccsu-2A.1*), TraesCS4A02G008000 (*QSd.ccsu-4A*), and TraesCS7A02G141300 (*QFf.ccsu-7A*) were functionally linked to grain weight, spike density, and floret fertility traits, respectively. Besides, 31 more CGs were involved in key physiological responses including oxidative stress, proline accumulation, membrane stability, leaf senescence, spikelet fertility, and grain weight (Supplementary Table S9).

Notably, 19 CGs underpinning 16 QTL for six spike traits encode protein kinases, central to signal transduction via phosphorylation—consistent with their known role in heat stress responses (Praat *et al.,* 2021; Sharma *et al.,* 2022). Several CGs encode proteins previously implicated in heat tolerance, including F-box, AP2/ERF, PPR, HSF-type, DNA-binding, NAC transcription factors, and ubiquitin-like domain proteins, as demonstrated in wheat, Arabidopsis, tobacco, and algae (Guo *et al.,* 2015; Li *et al.,* 2018; Xie *et al.,* 2019; Bi *et al.,* 2020; Pengyan *et al.,* 2020; Guo *et al.,* 2021). Overexpression studies confirm their functional roles, e.g., *TaFBA1* in Arabidopsis and tobacco (Li *et al.,* 2018), *TaNAC2L* (Guo *et al.,* 2015), and *TaHsfA6f* (Bi *et al.,* 2020). Additionally, RT-qPCR profiling of the AP2/ERF gene distinguished heat-tolerant from heat-sensitive wheat genotypes, reinforcing its regulatory significance in heat stress adaptation (Magar *et al.,* 2022).

Six QTL (*QGws.ccsu-7B*, *QGws.ccsu-7A*, *QNss.ccsu-7D*, *QNss.ccsu-6B.1*, *QSl.ccsu-6B*, and *QNfs.ccsu-1B*) co-located with key genes including *TaHsfC2a-B*, *TaCol-B5*, *WAPO1*, *Starch synthase I*, *TaGW2-B1*, and *PI1-1B/WPI-1-1B*—linked to NFS, NSS, SL, and GWS (Table 3; Supplementary Fig. S3). Among these, *TaHsfC2a-B* and *Starch synthase I* contribute to thermotolerance (Hu *et al.,* 2018; Tian *et al.,* 2018), while *WAPO1*, *PI1-1B/WPI-1-1B*, *TaGW2-B1*, and *TaCol-B5-7A* regulate spikelet number, spike development, grain weight, and spike architecture (Kuzay *et al.,* 2019; Schilling *et al.,* 2020; Zhang *et al.,* 2018, 2022). These associations underscore the functional relevance of the mapped QTL.

Despite their significance, these genes have yet to be utilized in MAS. Development of breeder-friendly KASP markers for these loci would facilitate their deployment in breeding. MAS is particularly effective when target genes reside in high-recombination telomeric regions, though even centromeric loci can be exploited via CRISPR-Cas9. For instance, *TaHsfA1* has been validated for heat tolerance in wheat using CRISPR-Cas9 (Wang *et al.,* 2023; Ikram *et al.,* 2024), highlighting the potential of genome editing in targeting heat-responsive genes.

### KASP and gene-based marker development

This study also focused on developing KASP and gene-based functional markers for stable QTL and key CGs associated with heat tolerance. A single KASP marker (KSNP8647_521) was successfully developed for floret fertility QTL *QFf.ccsu-3A* on chromosome 3A, detected under VLS conditions, and showed significant phenotypic segregation in the DH population (Fig. 4b). However, KASP marker development for the remaining 11 selected QTL was unsuccessful due to monomorphism or indistinct cluster separation—an outcome consistent with earlier wheat studies (Anuarbek *et al.,* 2019; Grewal *et al.,* 2022). In our recent work, only 3 of 14 targeted SNPs yielded successful KASP assays (Kumar *et al.,* 2024), highlighting the challenge, particularly when working with random SNPs from sequencing data versus gene-based markers (Kumar *et al.,* 2023). These findings emphasize the need for refined strategies in functional marker development for effective deployment in breeding programs.

In addition to the KASP marker, pangenome aided gene-based functional SSR markers were successfully generated for five of the eight CGs linked to spike traits and abiotic stress tolerance, with polymorphism observed among 16 wheat genotypes (Fig. 6b, d, e, f, h). Two SSR markers KAR3A (C)_n_ and CS6B (CT) were polymorphic between parental genotypes (PW343 and KSG1203) of the DH mapping population. One SSR showed segregation distortion, while the other fit the expected 1:1 ratio in the DH lines (Fig. 7). DH lines carrying the favorable allele for CS6B (CT) exhibited ∼1% higher FF, supporting its potential in breeding for terminal heat tolerance. Additionally, pangenome analysis also revealed SNPs and Indels within the eight CGs, offering further opportunities for gene-based marker development. These findings highlight the value of functional marker strategies leveraging structural variation and pangenome resources for trait-targeted wheat improvement.

## Abbreviations

IGP: Indo-Gangetic plains
DH: doubled haploid
PVE: percent variance explained
QTL: quantitative trait locus
SNP: single nucleotide polymorphism
TS: timely sown
LS: late sown
VLS: very late sown
FF: floret fertility
SL: spike length
NSS: number of spikelets per spike
SD: spike density
NFS: number of florets per spike
GWS: grain weight per spike
CIM: composite interval mapping
MAS: marker-assisted selection
MARS: marker-assisted recurrent selection
SBG: sequencing-based genotyping
KASP: kompetitive allele specific PCR
CG: candidate gene
CDS: coding DNA sequence
Indel: insertion and/or deletion
SSR: simple sequence repeat

## Supplementary data

The following supplementary data are available.

Fig. S1. Daily minimum and maximum temperature (°C) fluctuations at Lucknow location.

Fig. S2. Correlation among six spike related traits in each of the timely, late and very late sown environments.

Fig. S3. Distribution of 51 QTL for 6 spike related traits on the physical map of 18 wheat chromosomes.

Fig. S4. Heat maps representing differential expression of candidate genes underlying QTL.

Fig. S5. Knowledge network worked out by KnetMiner for seven important candidate genes.

Fig. S6. Structural variations (SNPs and Indels) across wheat cultivars based on CDS alignments in pangenome database.

Table S1. Procedure of phenotyping of six spike-related traits.

Table S2. Allele-specific forward primer and common primer sequences of KASP-SNPs.

Table S3. Forward (F) and reverse (R) primer sequences of five gene-based SSRs.

Table S4. List of 16 wheat accessions used for validation of SSR markers.

Table S5. Summary statistics including descriptive statistics, CV% (coefficient of variation), variance components and heritability estimates.

Table S6. Combined ANOVA showing mean squares for main and interaction sources of variation for the six spike traits.

Table S7. Path coefficient analysis showing direct (diagonal) and indirect (off-diagonal) effects.

Table S8. Detailed comparison of our study with the studies on meta-QTL (M-QTL) analysis for heat tolerance.

Table S9. High confidence candidate genes underlying spike-related QTLs for heat tolerance.

Table S10. Mining of SSRs across wheat genotypes used in pangenome development.

## Acknowledgements

The present study was supported by the Biotechnology Industry Research Assistance Council (BIRAC), Government of India, New Delhi and United States Agency for International Development Feed the Future Innovation Lab-Climate Resilient Wheat. Thanks are due to Dr. Anil Kumar Malik, Professor & Head, Department of Physics, Chaudhary Charan Singh University, Meerut for providing high performance computer facilities. HSB was awarded Honorary Scientist position by the Indian National Science Academy (INSA), New Delhi during the course of this study. The Heads, Department of Genetics and Plant Breeding, and Department of Botany and Chaudhary Charan Singh University, Meerut provided necessary facilities for this study. We also thank Dr. Anshu Chaudhary, Department of Zoology, Chaudhary Charan Singh University, Meerut for providing support to perform KASP genotyping.

## Author contributions

SK, HSB and KSG: conceptualization, design, and wrote the manuscript, with contributions from all the authors; SK, SoK, VPS, HS, RA, SKB and RV: conducted field experiments, trait phenotyping, genetic analysis and marker development; KSR and KSG: performed sequencing based genotyping; SK and SoK: interpreted the results and prepared first draft of the manuscript; SK, HSB and KSG: critically revised and edited the manuscript. All authors contributed to the manuscript and approved the submitted version.

## Conflict of interest

The authors of this manuscript declare no competing interest.

## Funding

This research was supported by a grant (No. BIRAC/TG/USAID/08/2014) from the Biotechnology Industry Research Assistance Council (BIRAC), Government of India, New Delhi, and the United States Agency for International Development (USAID) Feed the Future Innovation Lab-Climate Resilient Wheat.

## Data availability

Relevant data are included and referenced in this paper and its associated supplementary data.

## Notes

### Competing Interest Statement

The authors have declared no competing interest.

